# ChEC-seq: a robust method to identify protein-DNA interactions genome-wide

**DOI:** 10.1101/2021.02.18.431798

**Authors:** Maria Jessica Bruzzone, Benjamin Albert, Lukas Hafner, Slawomir Kubik, Aleksandra Lezaja, Stefano Mattarocci, David Shore

**Affiliations:** Department of Molecular Biology and Institute of Genetics and Genomics of Geneva (iGE3), University of Geneva, 30 quai Ernest-Ansermet, 1211 Geneva 4, Switzerland

## Abstract

Mittal et al. (2021; first brought to our attention in May 2019) have raised concerns regarding the Chromatin Endogenous Cleavage-sequencing (ChEC-seq) technique (Zentner et al., 2015) that may create a false impression that this method has fundamental flaws which prevent one from distinguishing between signal and noise. Although Mittal et al. focus on studies of the global co-activators SAGA, TFIID and Mediator that we were not involved in, we feel obliged to highlight here several of our own publications (Albert et al., 2019; Bruzzone et al., 2018; Hafner et al., 2018; Kubik et al., 2019; Kubik et al., 2018), as well as recent unpublished data, that employed ChEC-seq and directly addressed the observation raised by Mittal et al. that cleavage maps for various MNase fusion proteins often qualitatively resemble each other and those generated by “free” (unfused) MNase. Our studies lay out a clear path for determining sites of preferential factor localization by normalization of ChEC-seq experimental data to matched free-MNase controls. They also demonstrate the use of in vivo functional assays to assess ChEC-seq reliability and reveal examples where ChEC-seq identifies functional binding sites missed by conventional ChIP-seq analysis.

---

We begin by discussing an analysis of the only essential chromatin remodeler in yeast, RSC (Kubik et al., 2018). Given that ChIP of chromatin remodelers is notoriously inefficient (Yen et al., 2012), we used ChEC-seq to ask where the essential RSC remodeler is bound genome-wide, comparing Rsc8-MNase read-count peaks to those of a free MNase control. Despite the qualitative resemblance of the experimental and control data sets, we found that Rsc8 ChIP-seq peaks are often higher than free-MNase peaks in comparisons of data sets with matched basal read-count levels (cf. Figure 3B; (Kubik et al., 2018)). Parenthetically, we have consistently noted that background matching typically requires a 20-minute digestion for cells expressing unfused MNase, approximately 5- to 20-fold longer than for various MNase fusions. The reason for this is unknown but may be due to differences in nuclear concentrations, non-specific chromatin association, or some other factor(s).

To evaluate Rsc8 binding specificity we calculated the ratio of read counts at each peak to those at the same site in the control (free MNase) data set. To independently assess the validity of this approach we measured the effect of rapid nuclear depletion of RSC on nucleosome positioning genome wide, using the anchor-away technique and MNase-seq, respectively (Haruki et al., 2008; Kubik et al., 2018). This revealed a significant correlation between Rsc8-MNase read counts under peaks (both raw and normalized) and nucleosome occupancy changes in these regions (R=0.34; cf. Figure S3A, B in (Kubik et al., 2018)). In contrast, the correlations for free-MNase ChEC-seq or Rsc8 ChIP-seq (Yen et al., 2012) peaks were near zero (R=0.02 and -0.04, respectively; cf. Figure S3A, B in (Kubik et al., 2018)). Interestingly native ChIP-seq, which does not involve formaldehyde crosslinking and in this case was applied to the catalytic subunit of RSC, Sth1 (Ramachandran et al., 2015), correlated significantly better with our RSC depletion data (R= 0.21), though still less well than did ChEC-seq data. These comparisons suggest that, at least for the case of RSC, ChEC-seq is a reliable method for detection of functional interactions, particularly in comparison to standard ChIP-seq. A subsequent study from our lab indicates that the same is true for other chromatin remodelers (Kubik et al., 2019).

We now describe the case of Rif1 (Rap1-interacting factor 1; (Hardy et al., 1992)), a telomere-associated protein also implicated in temporal control of DNA replication origin firing ((Lian et al., 2011); reviewed in (Mattarocci et al., 2016)), which provides an instructive example of the power of ChEC-seq to identify what are likely to be multiple transient chromatin interactions. Since Rif1 was for many years undetectable at replication origins by ChIP, we decided to explore its chromatin association genome-wide by ChEC-seq. A quantitative analysis of the data revealed preferential binding near those replication origins whose firing is specifically affected by *RIF1* deletion (cf. Figure 3 in (Hafner et al., 2018)). Interestingly, although a more recent ChIP-seq analysis of Rif1 did reveal origin binding (Hiraga et al., 2018), signal strength did not correlate with the sites where Rif1 inhibits origin firing.

As its name implies, Rif1 protein interacts strongly with arrays of Rap1 that are bound to telomeric TG repeats at chromosome ends (Shi et al., 2013). This telomeric binding of Rif1 is detected robustly by both ChIP and ChEC. However, since Rap1 also functions as a transcription factor (TF) at > 300 promoters, one might imagine that Rif1 is recruited at these sites too, contrary to the claim by Mittal et al. (Mittal et al., 2021) that Rif1 is not expected to be enriched at promoters. Indeed, we found strong evidence that Rif1 associates specifically with promoter regions that are bound by Rap1 (cf. Figures 3C, 3D and related text in (Hafner et al., 2018)). Nevertheless, our report (Hafner et al., 2018) was focused on Rif1’s association with origins of DNA replication. We thus present here a more detailed analysis of these data which provides clear evidence that Rif1 associates with promoters through its ability to bind Rap1. The Rap1-Rif1 interaction requires a short C-terminal motif in Rif1, referred to as the Rap1 binding module (RBM; (Shi et al., 2013)). We therefore examined the ChEC-seq profile of rif1^RBM^-MNase, which contains point mutations in the Rif1 RBM that strongly diminish Rap1 binding both in vitro and in vivo (Shi et al., 2013). **Figure 1** shows two different genome browser displays of ChEC-seq data (taken from (Hafner et al., 2018)) for Rif1-MNase, rif1^RBM^-MNase, and free MNase, together with ChIP-seq data showing strong Rap1 binding at three different ribosomal protein genes (*RPS14A, RPS9B*, and *RPL21A*). Significantly, the strong Rif1-MNase signals coinciding with all three promoter sites of Rap1 binding are completely abolished by the RBM mutation. As expected, Rif1’s specific association with two nearby replication origins (*ARS313* and *ARS314*) is unaffected by the RBM mutation and serves as an internal control for the loss of rif1^RBM^-MNase association at the Rap1-bound promoters. For a global analysis of the effect of the RBM mutation on Rif1 association with Rap1, as measured by ChEC-seq, the reader is referred to Hafner et al. ((Hafner et al., 2018); cf. Figure 4C and linked data sets). We note that Hiraga et al. (Hiraga et al., 2018) also detected Rif1 at some Rap1-bound promoters by ChIP-seq and showed that this signal was abolished by a large deletion of the Rif1 C-terminus, which includes the RBM.

**Figure 1.**
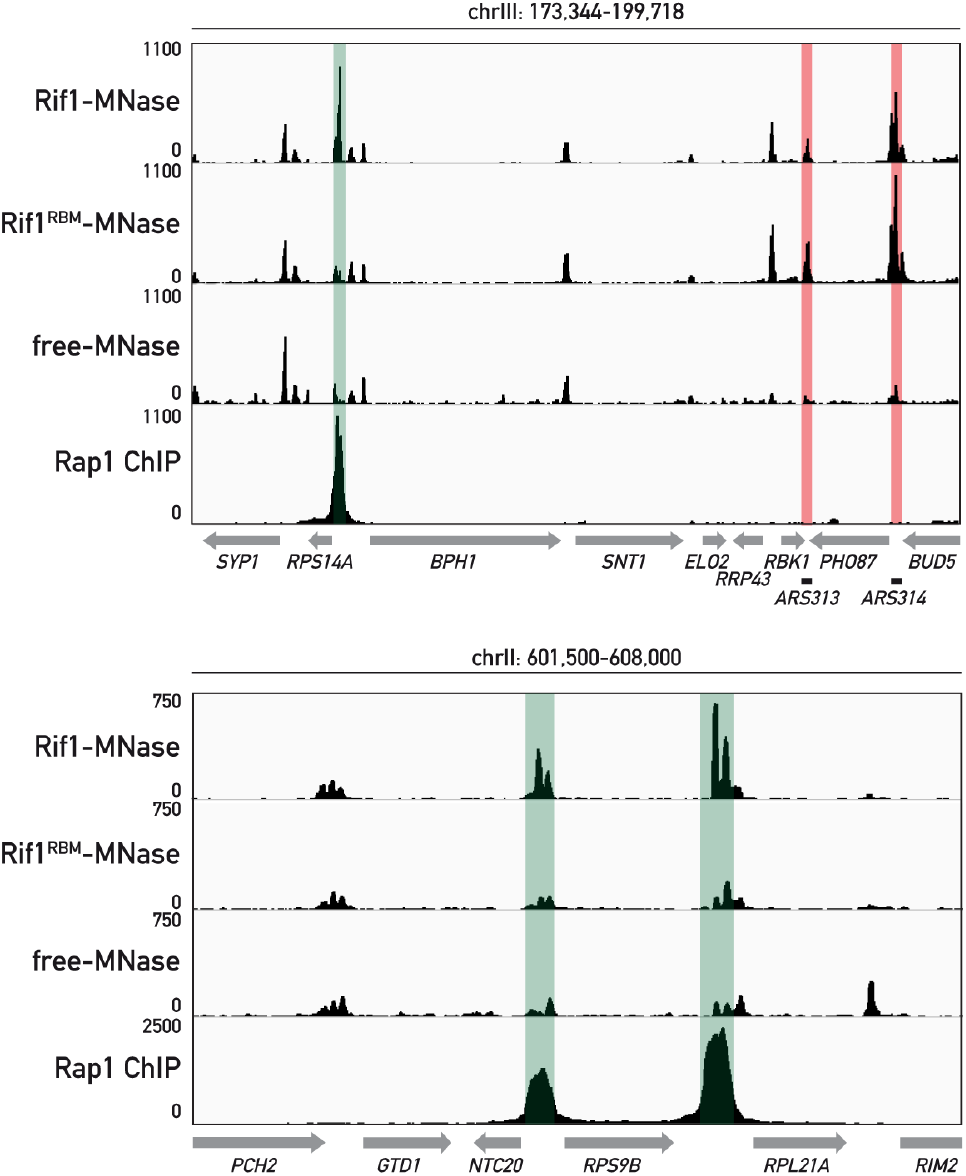
ChEC-seq reveals association of Rif1 with promoters bound by Rap1. The Rif1-MNase ChEC-seq signals coinciding with strong Rap1 binding at three different promoter regions (*RPS14A*, top panel, and both *RPS9A* and *RPL21A*, bottom panel; indicated by the green bars) is completely abrogated, i.e., reduced to free MNase background levels, when analyzed using the rif1^RBM^-MNase mutant protein in an otherwise identical ChEC-seq experiment. Note that the rif1^RBM^-MNase mutant shows wild-type levels of binding at two DNA replication origins (pink bars, top panel), as expected, and serves as an internal control.

Our analysis of Sfp1 (Split zinc-finger protein 1) provides a striking example where ChEC-seq reveals TF target sites that were undetectable by ChIP-seq (Albert et al., 2019). Transcriptome analysis of Sfp1 (Split zinc-finger protein 1) had suggested that it activates both ribosomal protein (RP) genes and a large suite of genes required for ribosome biogenesis (RiBi genes; (Cipollina et al., 2008)). Curiously though, ChIP assays were only able to detect Sfp1 at RP gene promoters and a few other genes, mostly with poor enrichment over background. Remarkably, ChEC-seq reveals strong Sfp1 binding at essentially all RiBi gene promoters, with read-count values strongly correlated with the strength of Sfp1-mediated activation (Albert et al., 2019). Interestingly, Sfp1 ChEC-seq peaks appear to reflect recognition of binding motif identified in vitro ((Zhu et al., 2009); cf. Figure 4C in (Albert et al., 2019)) which is similar to one found at many RiBi gene promoters (Hughes et al., 2000). To test directly the role of this motif _((A/G)(A/C)A_4_T_4_(C/T)), we have now mutated one such site_ at its chromosomal locus and tested the effect on Sfp1 binding by ChEC-seq. As shown in **Figure 2**, this 3 bp mutation at the bi-directional *RBD2-YPL245W* promoter region completely abrogates the Sfp1-MNase ChEC signal. Sfp1 binding at the nearby *SRP68* promoter is unaffected, as expected, and serves as an internal control for the specificity of the reduced signal at the *RBD2-YPL245W* promoter. These new ChEC-seq data provide the first direct evidence that the conserved motif identified in our study (Albert et al., 2019) is in fact an in vivo binding site for Sfp1 that has gone undetected by ChIP. As we noted previously (Albert et al., 2019), the failure to identify Sfp1 by ChIP at the promoters of RiBi and other growth-related genes might be due to the near absence of G/C base pairs in the binding motif, which may render its direct formaldehyde crosslinking to DNA extremely inefficient (Rossi et al., 2018). In contrast, ChIP-detected Sfp1 binding occurs at promoters that do not contain the Sfp1 binding motif and requires other TFs (e.g., Ifh1 at RP genes and Swi4 at G1/S regulon genes; (Albert et al., 2019)). We have thus proposed that ChIP-detected binding of Sfp1 may be indirect and largely independent of sequence-specific DNA interactions (Albert et al., 2019).

**Figure 2.**
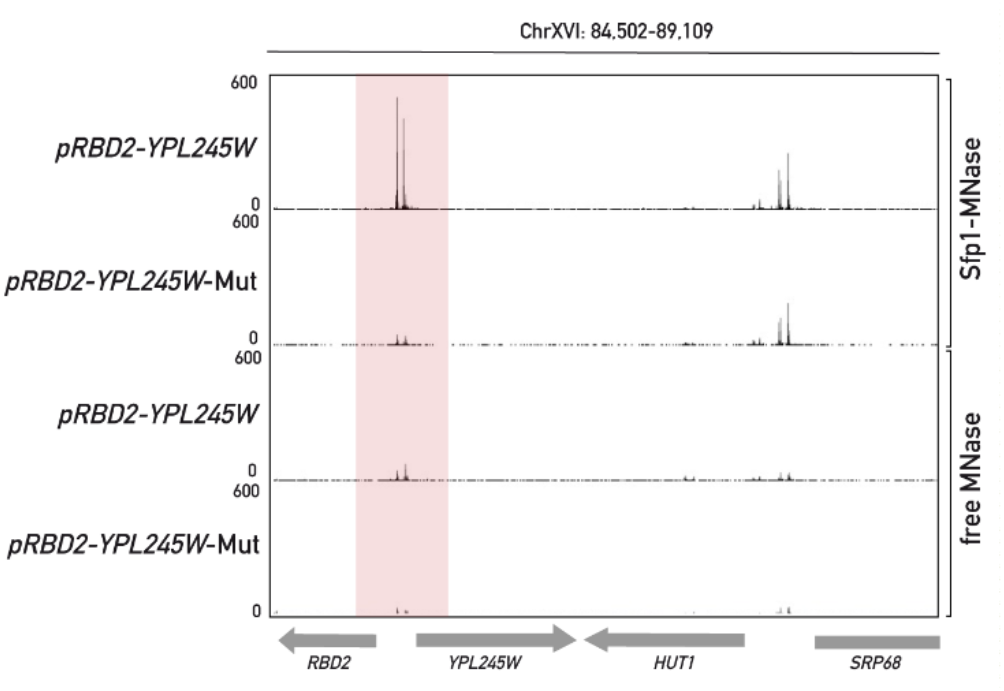
Promoter binding of Sfp1 at many genes involved in ribosome biogenesis and growth is dependent upon a specific A/T-rich DNA motif. Sfp1-MNase ChEC-seq signal at the bidirectional *RBD2-YPL245W* promoter (*pRBD2-YPL245W*; pink bar) is abolished by a triple point mutation in the conserved Sfp1 binding motif (TGAAAAATTTTC to TGAAGACTTCTC). Sfp1-MNase signal at a nearby promoter region, which directs expression of the *SRP68* gene, is unaffected, as expected, and serves as an internal control for the signal loss at *pRBD2-YPL245W*. Anchor-away studies have shown that both *RBD2, YPL245W*, and *SRP68* (but not *HUT1*) are activated by Sfp1 (Albert et al., 2019).

**Figure 3.**
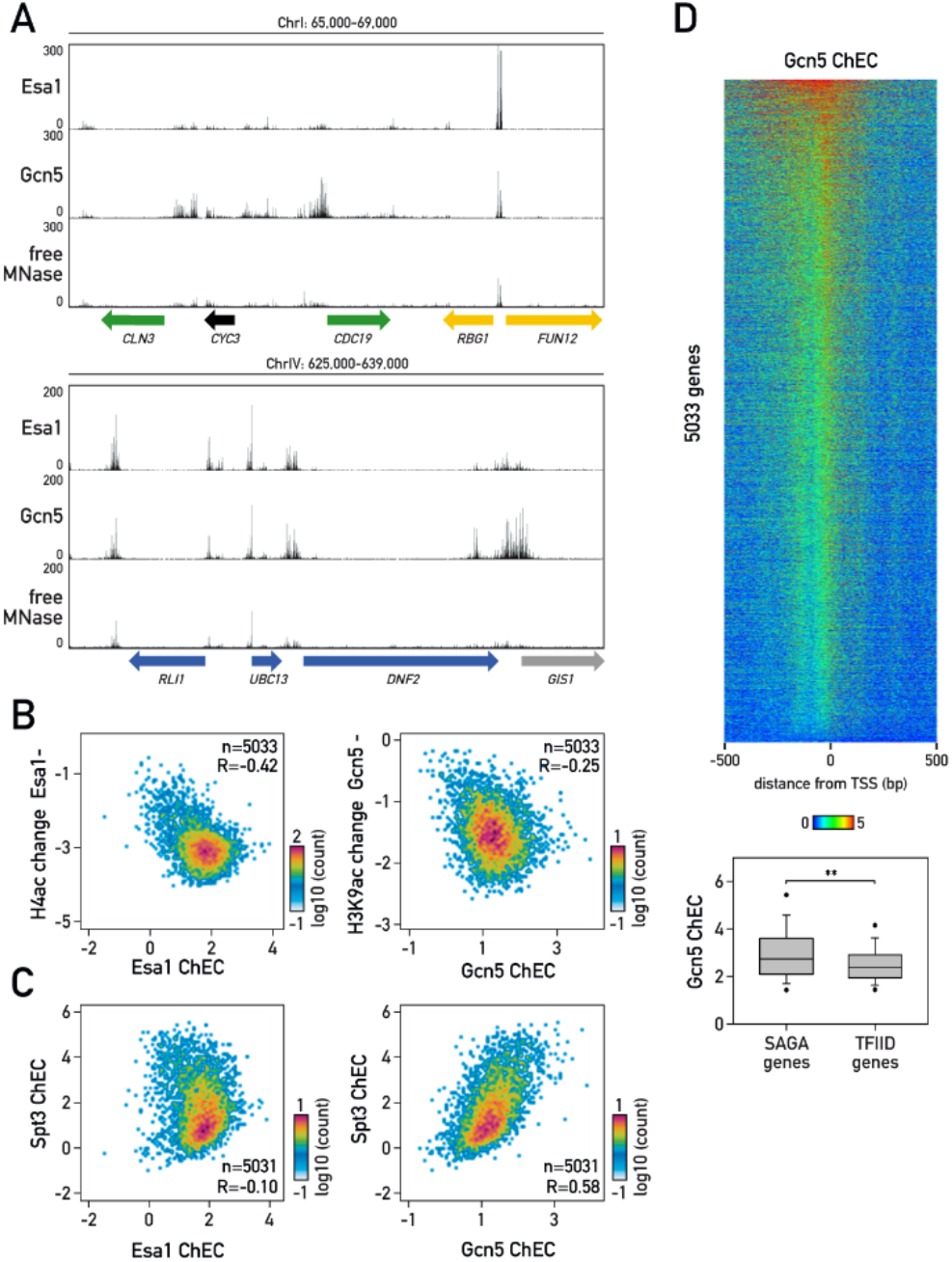
ChEC-seq reveals specific Esa1 and Gcn5 binding profiles. **(A)** Snapshots of ChEC-seq signal (read counts) for Esa1-MNase, Gcn5-MNase and a free MNase control at two representative genomic regions. For Esa1-MNase and Gcn5-MNase, samples were treated with 5 mM Ca+2 for 1 minute. ChEC-Seq was performed and analyzed as described in (Kubik et al., 2018). The free MNase control (5 mM Ca+2 treatment for 20 minutes) is from (Hafner et al., 2018). Genes are colored according to their Esa1 or Gcn5 transcriptional dependence, as measured by RNAPII ChIP-seq following anchor-away nuclear depletion (data from (Bruzzone et al., 2018)). Genes whose transcription is more dependent on Esa1 (yellow), more dependent on Gcn5 (green), equally dependent on both (blue), or dependent upon neither (black) are indicated at the bottom of each panel. RNAPII occupancy over the ORF of the SAGA-dominated gene *GIS1* (as defined in (Huisinga and Pugh, 2004)) is below background (Bruzzone et al., 2018) and thus its Esa1 or Gcn5 dependence cannot be determined. **(B)** The scatterplot on the left shows the relationship between histone H4 acetylation change at promoters following Esa1 nuclear depletion (Esa1-; from (Bruzzone et al., 2018)) and Esa1 ChEC-seq signal (normalized to a free-MNase control) at promoters for all protein-coding genes with an annotated transcription start site (TSS). The scatterplot on the right shows the relationship between histone H3K9 acetylation change at promoters following Gcn5 nuclear depletion (Gcn5-; from (Bruzzone et al., 2018)) and Gcn5 ChEC-seq signal (normalized to a free-MNase control) at the same set of RNAPII promoters. Normalized ChEC-seq signal for Esa1 and Gcn5 were measured in a 400 bp window upstream of the TSS. Axes on the scatterplots are in log2 scale. Sample size (n) and Pearson coefficient (R) are indicated. **(C)** Scatterplots showing relationship between Spt3 ChEC-seq signal (Baptista et al., 2017) and Esa1 ChEC signal (left) or Gcn5 ChEC-seq signal (right) at promoters for all protein-coding genes with an annotated TSS. ChEC-seq signal measurement and display are as in (B). **(D)** Heatmap of Gcn5 ChEC-seq signal normalized to free-MNase control around the TSS for all the yeast genes with an annotated TSS sorted according to the strength of the signal (top panel). Box plot (bottom panel) comparing Gcn5 ChEC-seq signal (normalized to free-MNase control) at promoters of SAGA-dominated (n=455) and TFIID-dominated (n=4219), as previously defined (Huisinga and Pugh, 2004). Asterisks indicate significant difference (p<0.01, Mann-Whitney test).

Finally, we present previously unpublished ChIP-seq data for Esa1, the catalytic subunit of the NuA4 histone acetyltransferase complex, and for the corresponding enzyme in SAGA, Gcn5. Nuclear depletion of Esa1 leads to a >1.5-fold reduction in RNA polymerase II (RNAPII) binding at about half of all protein-coding genes (Bruzzone et al., 2018). Consistent with this, Esa1 ChEC-seq reveals cleavage at many intergenic regions, qualitatively similar to that of SAGA components (Baptista et al., 2017). Nevertheless, we observed clear differences between the two, with Gcn5-dependent promoters displaying higher Gcn5 signal compared to Esa1 and vice versa (**Figure 3A**). Significantly, normalized ChEC-seq read-counts for both Esa1 and Gcn5 peaks correlate well with the sites of action of the corresponding enzymes as measured by acetylation-specific histone ChIP-seq (**Figure 3B**). Furthermore, and in contrast to the claim by Mittal et al., normalization of Gcn5 peaks to the free-MNase control does indeed indicate that SAGA binding is above background at many promoters, which may be obscured by comparing average read count values over thousands of promoters (cf. Figure 3 in Mittal et al.). The significance of Gcn5 and Esa1 ChEC-seq differences is further supported by the observation that Esa1 ChEC signals are uncorrelated (R=-0.10) with those reported for the SAGA component Spt3 (Baptista et al., 2017), whereas signals from two different SAGA subunits (Spt3 and Gcn5) are themselves strongly correlated (R=0.58; **Figure 3C**). We would also emphasize that normalized Gcn5 ChEC-seq signal strength is indeed strongest at promoters originally identified as SAGA-dependent ((Huisinga and Pugh, 2004); **Figure 3D**). Amongst the top 200 target genes, 26.5% are SAGA-dominated compared to 9% of the 5033 genes analyzed (p=5.25e-18, chi-square test). Finally, as a word of caution, we note that at least some of the Gcn5 targets that we identify are stress-induced genes, not expected to be active under the growth conditions used. Gcn5 binding in these cases may result from stress induced by the ChEC protocol itself, a point we plan to elaborate on elsewhere.

To summarize, we have demonstrated in several previous publications and by new data presented here that reliable quantitative information on the genome-wide binding of a diverse set of chromatin-associated factors can be derived from ChEC-seq experiments when read-count peaks are normalized to those of a free MNase control. These findings, backed up by independent functional assays, argue that ChEC-seq is a robust and sensitive method for identifying binding sites of chromatin-associated factors.

In closing, we would like to note, as mentioned above, that the issues raised by Mittal et al. (Mittal et al., 2021) focused primarily on ChEC-seq studies of the global co-activators SAGA, TFIID and Mediator. Their specific concerns are addressed in two separate reports from the authors of these studies (Donczew et al., 2021; Zentner et al., 2021). Finally, we would like to point the reader to a remarkable recent study (Brodsky et al., 2020) that demonstrates the power of ChEC-seq to measure subtle differences in the in vivo localization of groups of highly related transcription factors. We thus imagine that future applications of the ChEC-seq method will have important implications for the study of a wide range of problems related to chromosome biology.

## Material and Methods

### Yeast strains

*Saccharomyces cerevisiae* strains used in this study, all derived from W303-1**a** or W303-1α (Thomas and Rothstein, 1989) are listed in Supplemental Table 1. A triple point mutation at the *RPD2-YPL245W* promoter was introduced using the delitto perfetto method (Stuckey et al., 2011).

### ChEC-seq

Overnight yeast cultures were diluted to OD_600_=0.1 and grown in YPAD at 30°C. At OD_600_∼0.6, cells were washed and resuspended in a buffer containing 15 mM Tris, pH 7.5, 80 mM KCl, 0.1 mM EGTA, 0.2 mM spermine, 0.5 mM spermidine, 1× Roche EDTA-free mini protease inhibitors, 1 mM PMSF and 0.1% digitonin and incubated for 5 min at 30°C. CaCl_2_ was added to the final concentration of 5 mM and reactions were stopped after 1 (for MNase-fused proteins) or 20 minutes (for free MNase) by adding EGTA to a final concentration of 50mM. DNA was isolated and libraries were prepared as previously described (Hafner et al., 2018; Kubik et al., 2019). Libraries were sequenced using a HiSeq 2500 in single-end mode. Reads were mapped to the *S. cerevisiae* genome (sacCer3 assembly) using Bowtie2 through HTSStation (David et al., 2014). Read densities were normalized to 10M reads. For each read the 5’-most base was taken as the position of the MNase cut site.

## Data availability

The data are available at GEO under accession number GSE133645.

## Acknowledgments

We thank members of the Shore lab for comments and discussions throughout the course of this work; Mylène Docquier and the Genomics Platform of iGE3 at the University of Geneva (https://ige3.genomics.unige.ch/) for high-throughput sequencing services; and Nicolas Roggli for his expert assistance with data presentation and manuscript preparation. D.S. acknowledges funding from the Swiss National Science Foundation (grant number 31003A_170153) and the Republic and Canton of Geneva.

