## Supplemental Table S1 for "ChEC-seq: a robust method to identify protein-DNA interactions genome-wide"

| Strain | Genotype* | Parent | Reference |
|---|---|---|---|
| HHY168 | *MATα tor1-1 fpr1::NAT RPL13A-2XFKB12::TRP1* | | (Haruki et al., 2008) |
| YJB475 | *MATα tor1-1 fpr1::NAT RPL13A-2XFKB12::TRP1 GCN5-3XFLAG-MNase::HIS3* | HHY168 | This study |
| YJB478 | *MATα tor1-1 fpr1::NAT RPL13A-2XFKB12::TRP1 ESA1-3XFLAG-MNase::HIS3* | HHY168 | This study |
| YDS2 (W303-1**a**) | *MAT***a** *leu2-3,112 his3-11,15 ura3-1 ade2-1 trp1-1 can1-100* | | (Thomas and Rothstein, 1989) |
| YDS3 (W303-1α) | *MATα leu2-3,112 his3-11,15 ura3-1 ade2-1 trp1-1 can1-100* | | (Thomas and Rothstein, 1989) |
| YJB564 | *MATα* p*YPL245W*-mut *SFP1-3XFLAG-MNase::hphMX4* | YDS3 | This study |
| YJB566 | *MATα* p*GUA1*-mut *SFP1-3XFLAG-MNase::hphMX4* | YDS3 | This study |
| YJB571 | *MATα* p*YPL245*-mut *pREB1-3XFLAG-MNase::URA3* | YDS3 | This study |
| YJB572 | *MAT***a** p*GUA1*-mut p*REB1-3XFLAG-MNase::URA3* | YDS2 | This study |

*All strains are of the W303 background (*leu2-3,112 his3-11,15 ura3-1 ade2-1 trp1-1 can1-100*; (Thomas and Rothstein, 1989)). Only additional modifications and the allele at *MAT* are indicated.
